# Evolutionary barriers to horizontal gene transfer in macrophage associated *Salmonella*

**DOI:** 10.1101/2022.04.01.486712

**Authors:** Rama P. Bhatia, Hande A. Kirit, Cecil M. Lewis, Krithivasan Sankaranarayanan, Jonathan P. Bollback

## Abstract

Horizontal gene transfer (HGT) is a powerful evolutionary force facilitating bacterial adaptation and emergence of novel phenotypes. Several factors, including environmental ones, are predicted to restrict HGT, but we lack systematic and experimental data supporting these predictions. Here, we address this gap by measuring the relative fitness of genes horizontally transferred from E. *coli* to *S. enterica* in infection relevant environments. We estimated the distribution of fitness effects in each environment and identified dosage-dependent effects across different environments are a significant barrier to HGT. The majority of genes were found to be deleterious. We also found longer genes had stronger negative fitness consequences than shorter ones, showing that gene length can significantly hinder HGT. Furthermore, fitness effects of transferred genes were found to be environmentally dependent. In summary, a substantial fraction of transferred genes had a significant fitness cost on the recipient, with both gene characteristics and the environment acting as evolutionary barriers to HGT.

## Introduction

Horizontal gene transfer (HGT) is the lateral transfer of genetic material between individuals of the same or different species rather than vertically from parent to offspring. HGT plays an important role in bacterial adaptation: e.g., acquisition of accessory genes can confer bacteria with novel phenotypes such as antimicrobial resistance, microbial toxins, and tolerance to heavy metals ^1^.

Gene acquisition through HGT may exert different fitness effects on the recipient cell, with the nature of these effects determining the evolutionary fate of the newly transferred gene. In the simplest terms, fitness can be defined as reproductive success and is key to understanding bacterial evolution and adaptation ^2^. A fitness advantage or disadvantage can be considered at different levels of the biological hierarchy (e.g., genes, transposons, plasmids, cells) that are under selection ^3,4,5,6^

Nonetheless, the sole purpose of this study was to explore which of the intrinsic properties or features of a horizontally transferred gene, upon its expression, could impact the fitness/reproductive ability of the host and not to understand fitness costs due to physical barriers to the acquisition and integration (e.g., success of transmission or probability of homologous or illegitimate recombination) of the transferred genes. In this study, the transferred genes which on expression significantly increased the relative fitness of the host were said to have a beneficial effect; those that caused a significant decrease in the relative fitness were said to have a deleterious fitness effect and those that did not significantly differ from the ‘wild type’ were classed as neutral.

While genes with a beneficial or neutral effect can persist in bacterial populations, genes with deleterious effects will most likely be purged ^7,8^. The relative frequency of these fitness effects, also known as the distribution of fitness effects (DFEs) ^9,10^ is thus an important predictor of the outcome of HGT events, as it provides insights into the evolutionary fate of a horizontally transferred gene determined by its fitness on the recipient cell. Therefore, understanding the role of factors in defining the DFEs of horizontally transferred genes is critical if we wish to understand bacterial evolution. Collectively, we will refer to these factors as evolutionary barriers to HGT.

Several potential evolutionary barriers to HGT have been predicted using computational approaches like phylogenetic and parametric/compositional analysis ^11^, including those due to *(i)* gene function: information processing, such as those involved in DNA replication, transcription and translation are less often transferred compared to operational genes ^12,13,14^, *(ii)* gene interactions: genes with multiple interacting partners (e.g., protein-protein, metabolic or regulatory) are less likely to experience a successful gene transfer event ^15,16^, *(iii)* deviation in GC content and codon usage: HGT frequencies can be affected by the donor-recipient similarity barrier. Genes acquired from donors of dissimilar GC content and codon usage are more likely to be targeted by anti-HGT systems (e.g., R-M systems, Cas/CRISPR, H-NS) ^17,18,19,20,21^, and *(iv)* gene dosage/protein dosage: an increase in protein levels due to extra copies of an ortholog can disrupt the stoichiometric balance in the cell ^22^.

In addition to the intrinsic characteristics of the genes themselves, the environment can substantially alter the fitness effects of horizontally acquired genes. For example, genes that confer antibiotic resistance are beneficial in the presences of the antibiotic but might be deleterious in its absence ^23,24^. In fact, multiple studies have shown that the environment may affect the selective outcomes in evolving bacterial populations ^25,26,27^.

Here, we experimentally test the fitness effects of HGT events, by introducing genes Escherichia coli into *Salmonella enterica* serovar Typhimurium strain 4/74 (*S*. Typhimurium 4/74) in four different infection relevant environments. *S*. Typhimurium is an enteric pathogen infecting both humans and animals and can cause non-typhoidal Salmonella disease (NTS), including a virulent invasive form of the disease in sub-Saharan Africa ^28,29,30^. The strain used in this study, *S*. Typhimurium 4/74, is a prototrophic bacteria that is highly virulent in cattle, pigs, mice, and chickens ^31,32^. *Salmonella* is an intracellular pathogen, and to survive and proliferate within eukaryotic cells, it must adapt to a fluctuating intracellular environment. During the course of evolution, *Salmonella* has acquired virulence encoded Salmonella Pathogenicity Islands (SPI) through horizontal gene transfer. Among many other pathogenic properties, SPIs have allowed *Salmonella* to hoodwink host immune surveillance and survive in a replicative niche within a modified phagosome in the host macrophages, known as the *Salmonella*-containing vacuole (SCV) ^28^.

We tested the consequences on fitness of the 44 transferred genes in environments that mimic key features of the macrophage environment ^32^; SPI-2 inducing PCN (phosphate/carbon/nitrogen) medium ^33,34^ as the base, and further modified it to either contain low magnesium, low oxygen (hypoxic), or an antibiotic challenge using a fluoroquinolone class of drugs – ciprofloxacin, a first-line antimicrobial for many Gram-negative pathogens, including *Salmonella* (see **Supplementary Table 1**).

In contrast to a retrospective computational approach, our experimental approach allows us to test which genes are likely to fail to survive the evolutionary sieve, and thus disentangle the influence of different evolutionary barriers and the role of the environment.

## Results

The experimental approach was to artificially transfer 44 orthologs from E. coli to Salmonella Typhimurium 4/74 using a low copy (3-4/cell) inducible expression vector pZS* ^35^ in four infection relevant environments (see **Supplementary Table 1**) to study the DFE of horizontally acquired genes. Additionally, we wanted to investigate the extent of the effects of evolutionary barriers – functional category, gene interactions, GC content, codon usage bias, gene length, protein dosage, and the environment on the persistence of horizontally transferred genes.

*S*. Typhimurium and *E. coli* were chosen primarily due to their genetic and ecological similarity, and history of HGT ^36,37^. Furthermore, the genetic similarity between the donor and recipient species ensures the functioning of the transferred genes in the host background, as well as that the functional interacting partners exist for the majority of genes transferred; this is important as the study aims to understand the role of gene categories and gene interactions in the persistence of horizontally acquired genes. Additionally, the donor and recipient are sufficiently divergent so that the effect of several intrinsic factors of the transferred genes can be evaluated, such as their GC content and length.

### DFEs and role of other barriers to HGT

Specifically, we looked at the effect of the functional category, gene interactions, GC content, codon usage bias, and gene length of the transferred orthologs on the fitness of *S*. Typhimurium 4/74 in four environments that mimic the signals encountered during an intracellular infection. We did this through careful and precise measurement of fitness effects from changes in gene frequencies using Next-Generation Sequencing (NGS). A graphical illustration of pooled growth setup has been provided **(Figure 1**).

**Figure 1.**
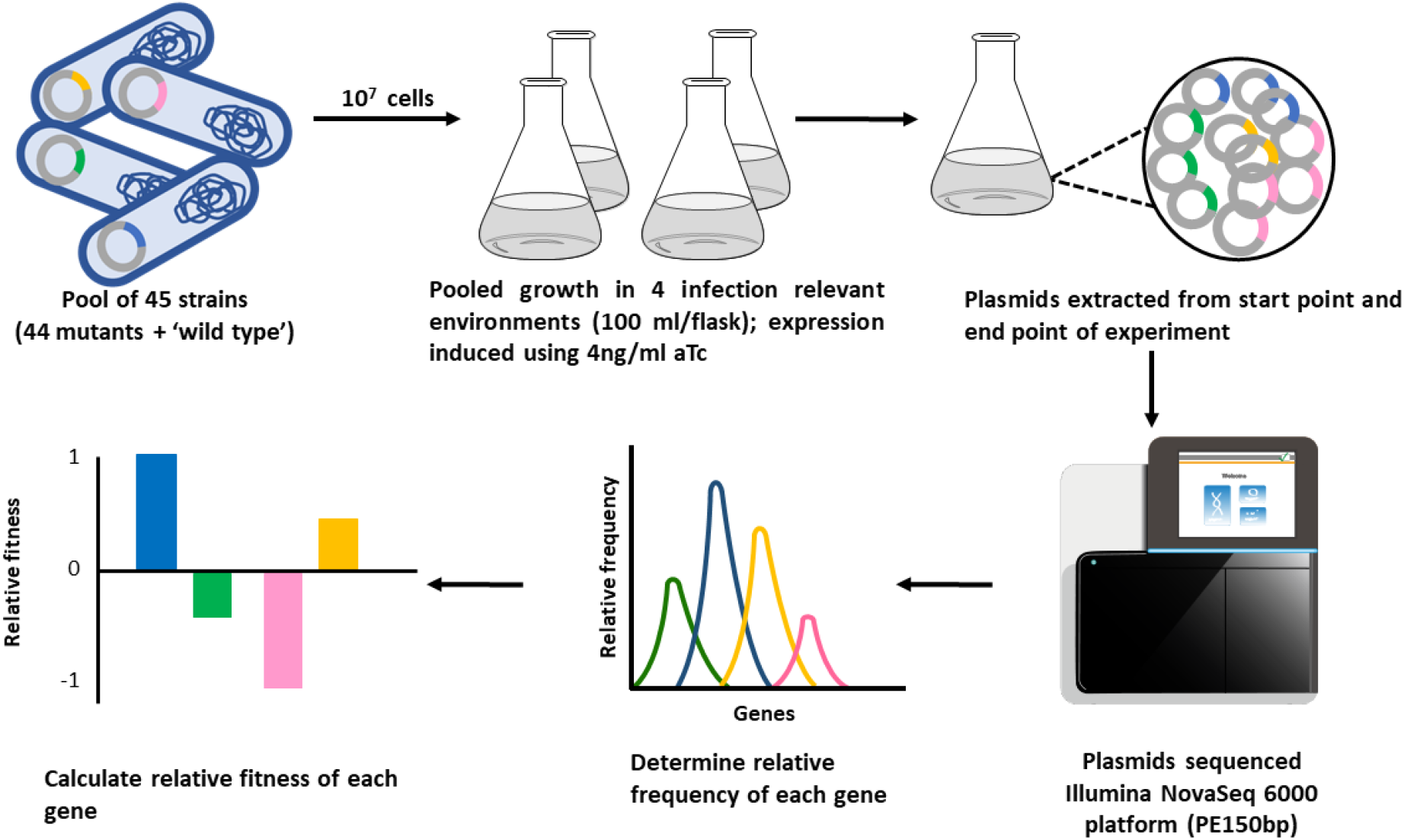
Graphical representation for pooled growth experiments. The flask-2 icon in the figure <u>by DCBLS 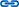</u> is licensed under <u>CC-BY 4.0 Unported 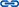</u>/ Object properties modified from original. The sequence_histogram icon by <u>Marcel Tisch 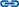</u> is licensed under <u>CC0 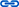</u> / Object properties modified from original. The genomesequencer9 icon by <u>DCBLS 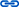</u> is licensed under CC-BY 4.0 <u>Unported 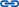</u>.

#### Estimating the distribution of fitness effects (DFEs) of horizontally transferred genes

The quantitative effects of mutations can be summarised as a distribution—the distribution of fitness effects, or DFE—which models the frequency of mutations with different strength of effects ^38^.

Many experimental studies have been carried out to determine the DFE of new mutations. To mention a few examples, DFEs have been estimated for *E. coli* mutants generated by random transposon mutagenesis ^39^, spontaneous and induced mutations in diploid yeast cell lines ^40^, for random mutations in vesicular stomatitis virus ^41^, for amino acid mutations in humans and *Drosophila melanogaster* ^10,42^, and for point mutations in genes ^43^. Parametric distributions like unimodal log-normal and gamma distributions, have been widely used to model and infer properties of DFEs ^44^. Although, there are differences in DFEs at the species and genomic level, they do exhibit some general properties, like beneficial/advantageous mutations are rare, strongly beneficial mutations are exponentially distributed and the DFE for deleterious/lethal mutations are complex and multi-modal ^38^.

In line with the previously observed DFEs ^41,45,46^, the fitness effects of our 44 genes in 4 different environments are best explained by a log-normal distribution (μ_InSPI2_ = −0.211, σ_InSPI2_ = 0.243 ; μ_InSPI2 Hypoxic_ = −0.174, σ_InSPI2 Hypoxic_ = 0.200 ; μ_InSPI2 Ciprofloxacin_ = −0.262, σ_InSPI2 Ciprofloxacin_ = 0.308 ; μ_InSPI2 Low Mg_ = −0.148, σ_InSPI2 Low Mg_ = 0.252) (Figure 2).

**Figure 2.**
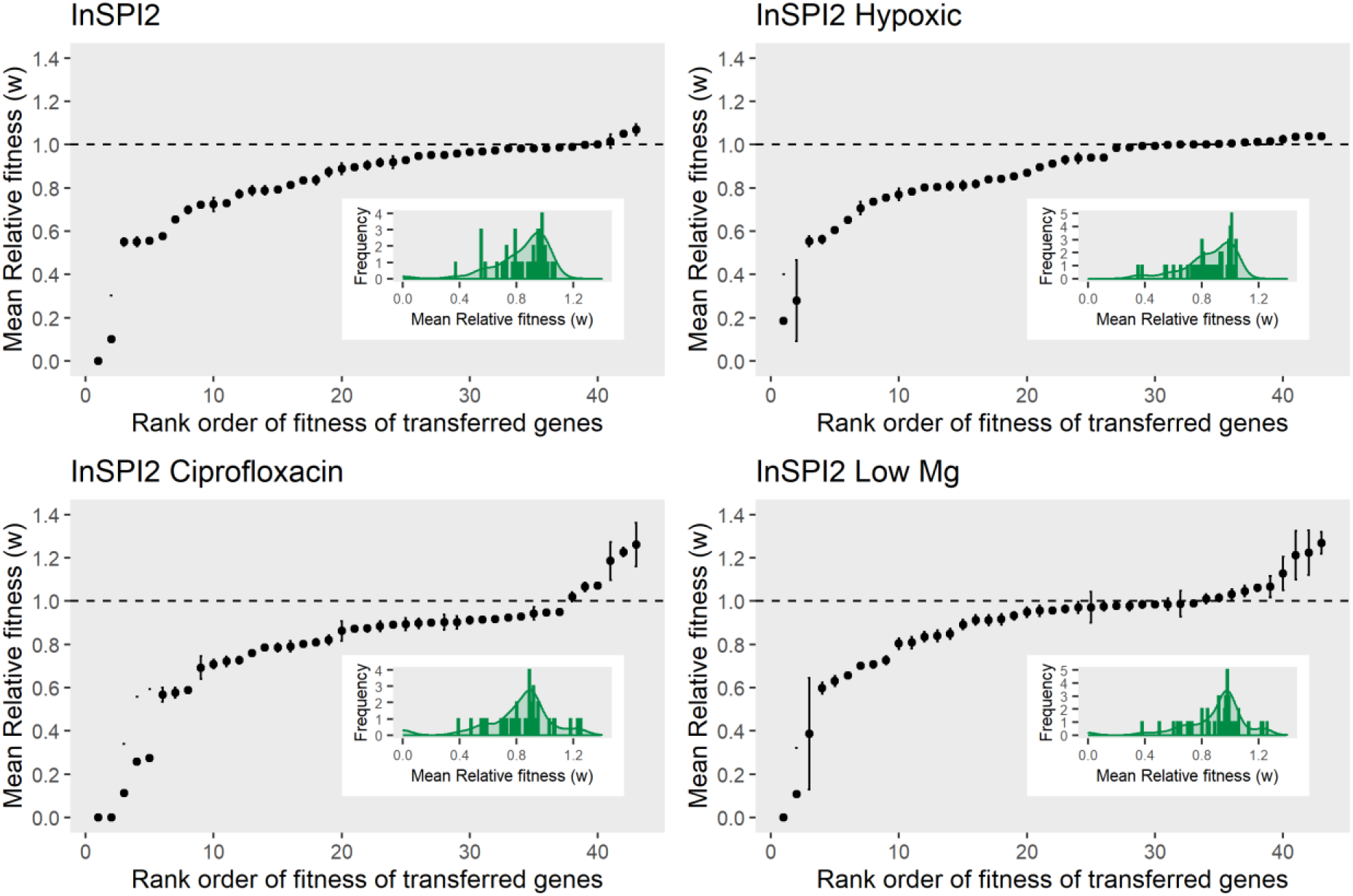
DFEs of transferred E.coli orthologs expressed in S. Typhimurium 4/74. The figure shows the distribution of fitness effects obtained from pooled growth assays in each infection relevant environment. On the x-axis, transferred genes are ordered by their relative fitness values in the respective environments. Error bars represent standard deviation of the mean for 4 replicate measurements for each gene. Embedded plots are histogram and density representations for the same data. Black dashed line shows a fitness of 1.

#### Categorising genes based on fitness effects

Based on the results from the one sample t-test, the genes were categorised into 5 groups in each environment with respect to their fitness effects (Figure 3). On average across the four environments ∼70% of the genes were deleterious, although ∼43% of these imposed a relatively minor fitness cost (mildly deleterious) (1 > mean fitness of gene > 0.8, P_*adj*_ < 0.05 with μ_0_ < 1), whereas ∼27% were highly deleterious (mean fitness of gene < 0.8, P_*adj*_ < 0.05 with μ_0_ < 1). Approximately 22% of the genes were found to be neutral (P_*adj*_ > 0.05 with μ_0_ > 1 and μ_0_ < 1) on average across the four environments (Figure 3). FDR corrected p-values for fitness effects of E.coli orthologs deviating from neutrality are provided as Supplementary Table 5.

**Figure 3.**
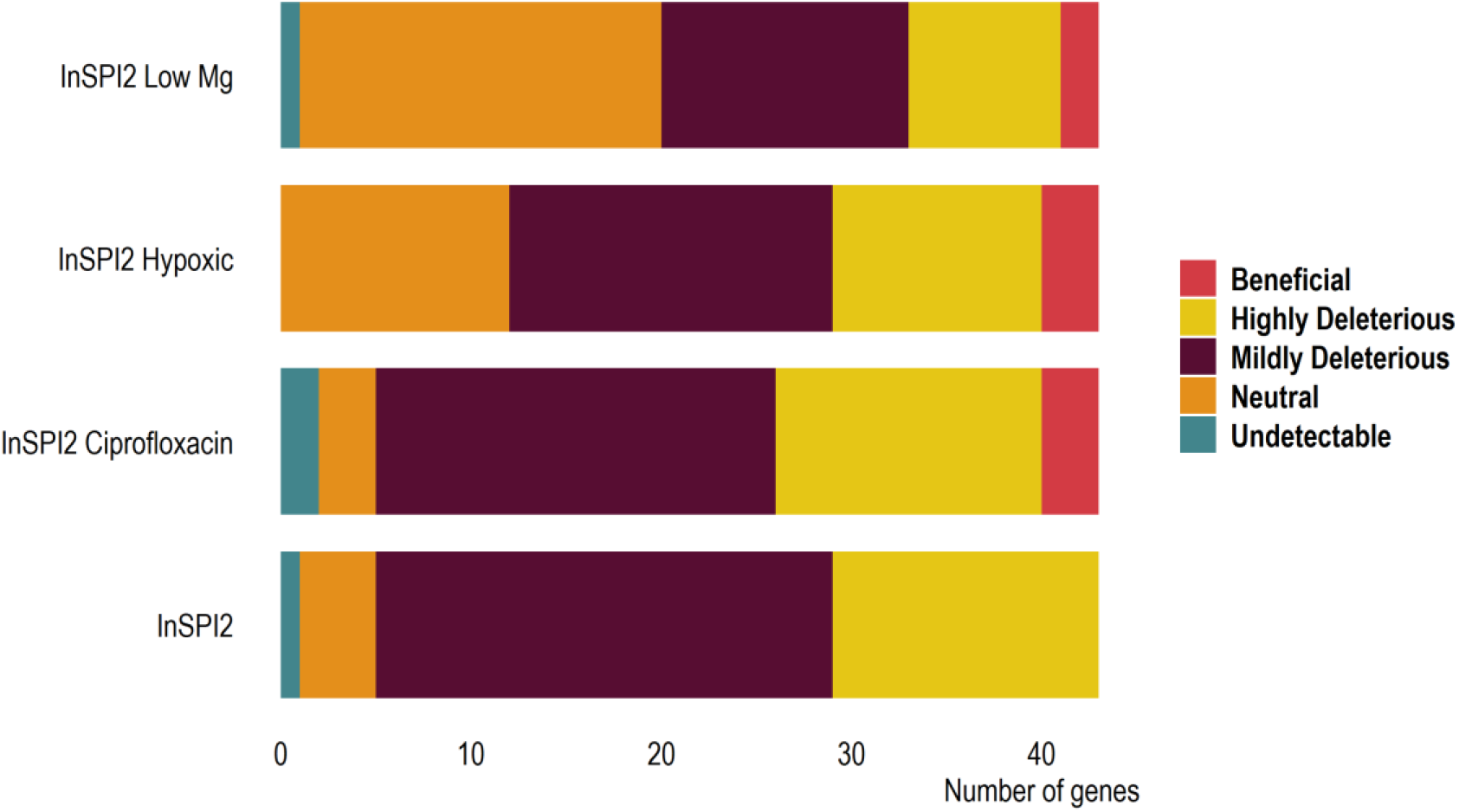
Grouping of genes based on fitness effects. Breakdown of the transferred orthologs based on results from one sample t-test in each growth environment. Undetectable genes: mean fitness of gene = 0, P_*adj*_ < 0.05 with μ_0_ < 1; Neutral genes: P_*adj*_ > 0.05 with μ_0_ > 1 and μ_0_ < 1; Mildly Deleterious genes: 1 > mean fitness of gene > 0.8, P_*adj*_ < 0.05 with μ_0_ < 1; Highly Deleterious genes: mean fitness of gene < 0.8, P_*adj*_ < 0.05 with μ_0_ < 1.; Beneficial genes: P_*adj*_ < 0.05, with μ_0_ > 1.

#### Functional category of genes

The ‘complexity hypothesis’ suggests that informational genes or genes involved in transcription, translation and replication are less likely to be horizontally transferred than the operational or housekeeping genes, those involved in amino acid synthesis, nucleotide biosynthesis, energy metabolism ^12,13,14^. We separated our transferred genes based on their COG annotations as informational (19 genes) and operational (20 genes) and tested whether or not the functional categories influenced the fitness effects of the transferred genes (excluding the genes that have not been classified (3 genes) and those that were classified as both informational and operational (2 genes)).

Contrary to the complexity hypothesis, we did not observe a significant difference in the mean fitness effects of genes between the two functional categories. This finding was consistent across all growth environments (InSPI2 : Med _Informational_ = 0.886, Med _Operational_ = 0.913, Wilcoxon rank sum test, W = 178, p = 0.482; InSPI2 Hypoxic : Med _Informational_ = 0.860, Med _Operational_ = 0.959, Wilcoxon rank sum test, W = 145, p = 0.158; InSPI2 Ciprofloxacin : Med _Informational_ = 0.890, Med _Operational_ = 0.884, Wilcoxon rank sum test, W = 192, p = 0.642; InSPI2 Low Mg : Med _Informational_ = 0.969, Med _Operational_ = 0.966, Wilcoxon rank sum test, W = 188, p = 0.597) (Figure 4). Although no significant difference was observed between the functional categories, genes with an average low fitness were mostly informational genes.

**Figure 4.**
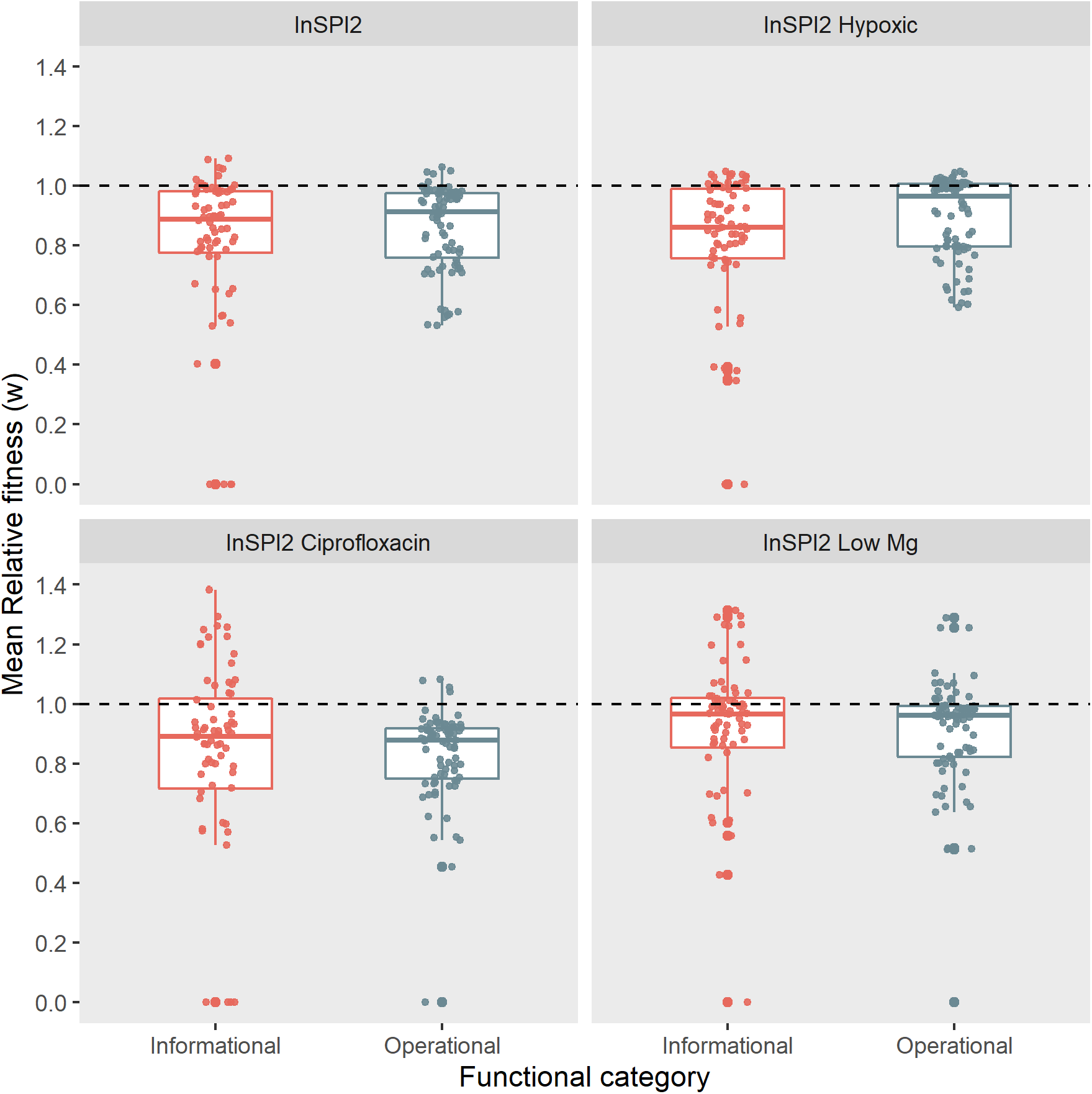
Fitness effects by functional categories. Boxplot with jitter points showing the fitness effects of transferred genes (4 replicate measurements) in four growth environments, with genes divided into two functional categories. Black dashed line shows a fitness of 1.

#### Complexity of gene interactions

The complexity hypothesis originally proposed that the transferability of genes was dependent on their biological functions ^13^, and later suggested that the factor affecting HGT was instead the number of connections in gene networks ^15^. The idea is that an increase in gene connectivity can reduce fitness of the bacterial host by creating a stoichiometric imbalance in the cell ^47,48^ or by improper interactions with their existing partners in the host ^22^.

To test whether high gene connectivity restricts HGT events, we investigated the relationship between fitness of the transferred orthologs and their corresponding interactions at three levels (protein-protein, metabolic, and regulatory) in *S*. Typhimurium 4/74. None of the gene interaction levels could significantly explain the fitness effects of the transferred orthologs in the four environments (Protein-protein interactions - InSPI2: R^2^ = 0.002, *p* = 0.75, InSPI2 Ciprofloxacin: R^2^ = 0.008, *p* = 0.75, InSPI2 Hypoxic: R^2^ = 0.004, *p* = 0.75, InSPI2 Low Mg: R^2^ = 0.002, *p* = 0.75; Regulatory interactions - InSPI2: R^2^ = 0.010, *p* = 0.70, InSPI2 Ciprofloxacin: R^2^ = 0.011, *p* = 0.70, InSPI2 Hypoxic: R^2^ = 0.002, *p* = 0.73, InSPI2 Low Mg: R^2^ = 0.009, *p* = 0.73; Metabolic interactions - InSPI2: R^2^ = 0.111, *p* = 0.05, InSPI2 Ciprofloxacin: R^2^ = 0.038, *p* = 0.20, InSPI2 Hypoxic: R^2^ = 0.09, *p* = 0.05, InSPI2 Low Mg: R^2^ = 0.097, *p* = 0.05) (Figure 5).

**Figure 5.**
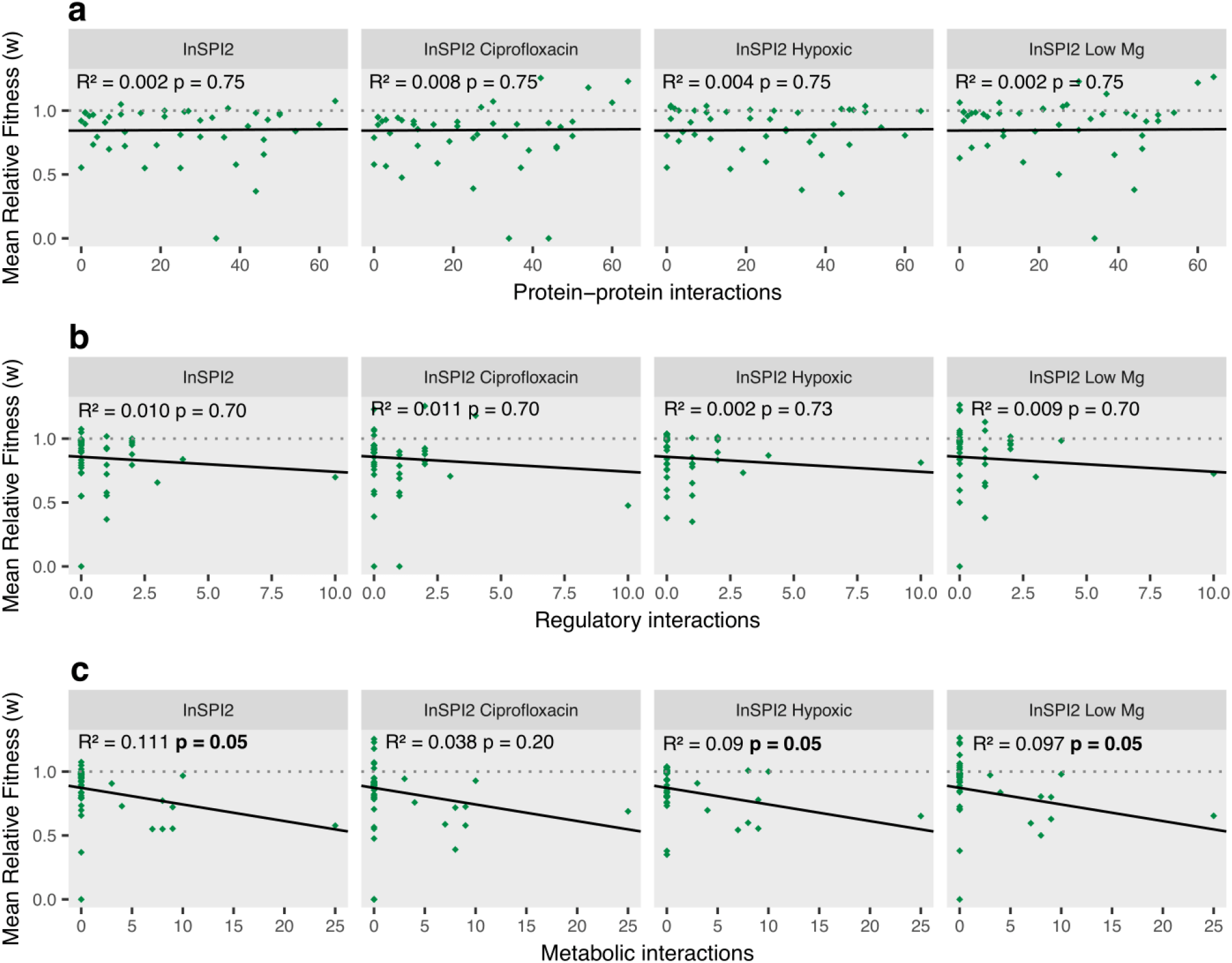
Relationship between relative fitness and number of interactions. Relative fitness of transferred E.coli orthologs in four growth environments plotted against the number of protein-protein interactions (**a**), regulatory interactions (**b**), and metabolic interactions (**c**) in S. Typhimurium 4/74. The black line is the regression between the two variables. Grey dotted line shows a fitness of 1. FDR corrected p-values and R^2^ values from the linear regression analysis are shown on the plot.

There might be a potential trend for increasing number of metabolic interactions leading to reduced fitness of *S*. Typhimurium 4/74 (Figure 5 c). However, based on a power analysis ^49^, the likelihood of the regression model to detect a significant trend was found to be 62%, suggesting that an increased number of high complexity genes might be needed to identify the effect of increasing metabolic interactions on HGT.

#### GC content and codon usage

The base composition similarity between ancestral genes within a genome aid in distinguishing a horizontally transferred foreign gene exhibiting different characteristics, like GC content and codon usage patterns, in comparison to the ancestral genes. It has been proposed that the horizontally acquired foreign genes gradually undergo an amelioration process, whereby the foreign genes shift their base composition and codon usage patterns to match the resident genome, thus making an HGT event undetectable through parametric computational analysis ^11,50^. Differences in the GC content of the foreign gene and recipient genome can impact the integration and persistence of the gene, and therefore can be a barrier to horizontal gene transfer. For example, the histone-like nucleoid structuring protein (H-NS) in *Salmonella* can silence the horizontally acquired foreign genes with a GC content lower than the recipient genome ^20^.

The acquired gene has a rich codon usage, if it has preferential usage for the abundant codons in the recipient genome, whereas if the gene has a bias for the rare codons, the gene is said to have a poor codon usage ^51^. Codon usage bias can affect the translational efficiency and accuracy of the cell ^52^. Incompatibility between the codon usage of the foreign gene and the recipient tRNA pool can lead to translational errors, for example, misfolded cytotoxic proteins. The misfolded proteins can cause aggregation at the cell membranes, thus disrupting the cell integrity and reducing the fitness of the cell ^53^.

The effects of GC content and codon usage on fitness were examined individually, although these measures are often highly correlated ^54^. We compared the absolute deviation in GC content between the transferred E. coli genes and their *S*. Typhimurium 4/74 orthologs with the observed fitness effects in four growth environments and did not find a significant relationship; InSPI2: R^2^ = 0.004, *p* = 0.79, InSPI2 Ciprofloxacin: R^2^ = 0.037, *p* = 0.79, InSPI2 Hypoxic: R^2^ = 0.005, *p* = 0.79, InSPI2 Low Mg: R^2^ = 0.00, *p* = 0.79 (**Supplementary Figure 3 a**). Similarly, the absolute deviation in the Frequency of Optimal codon usage (FOP) between the transferred *E. coli* genes and their *S*. Typhimurium 4/74 copies could not explain the observed fitness effects in any environment; InSPI2: R^2^ = 0.049, *p* = 0.29, InSPI2 Ciprofloxacin: R^2^ = 0.014, *p* = 0.29, InSPI2 Hypoxic: R^2^ = 0.056, *p* = 0.44, InSPI2 Low Mg: R^2^ = 0.022, *p* = 0.44 (**Supplementary Figure 3 b**).

#### Gene length is a significant predictor of HGT

The length of the gene can impose fitness costs at the chromosome, transcript, and protein levels in the host cell. Although, extra DNA is unlikely to be costly at the chromosome/replication level, the energy expenditure inflates with transcription and even more with protein synthesis ^55^. Ribosomal sequestration by lengthy genes will also limit the production of essential proteins, lowering host fitness and being selected against in a population ^56^. We observed a statistically significant negative relationship between gene length and the fitness effects of the transferred genes in all growth environments; InSPI2: R^2^ = 0.25, *p* = 6.17e-04, InSPI2 Ciprofloxacin: R^2^ = 0.13, *p* = 0.016, InSPI2 Hypoxic: R^2^ = 0.3, *p* = 1.29e-04, InSPI2 Low Mg: R^2^ = 0.13, *p* = 0.019) (**Figure 6**).

**Figure 6.**
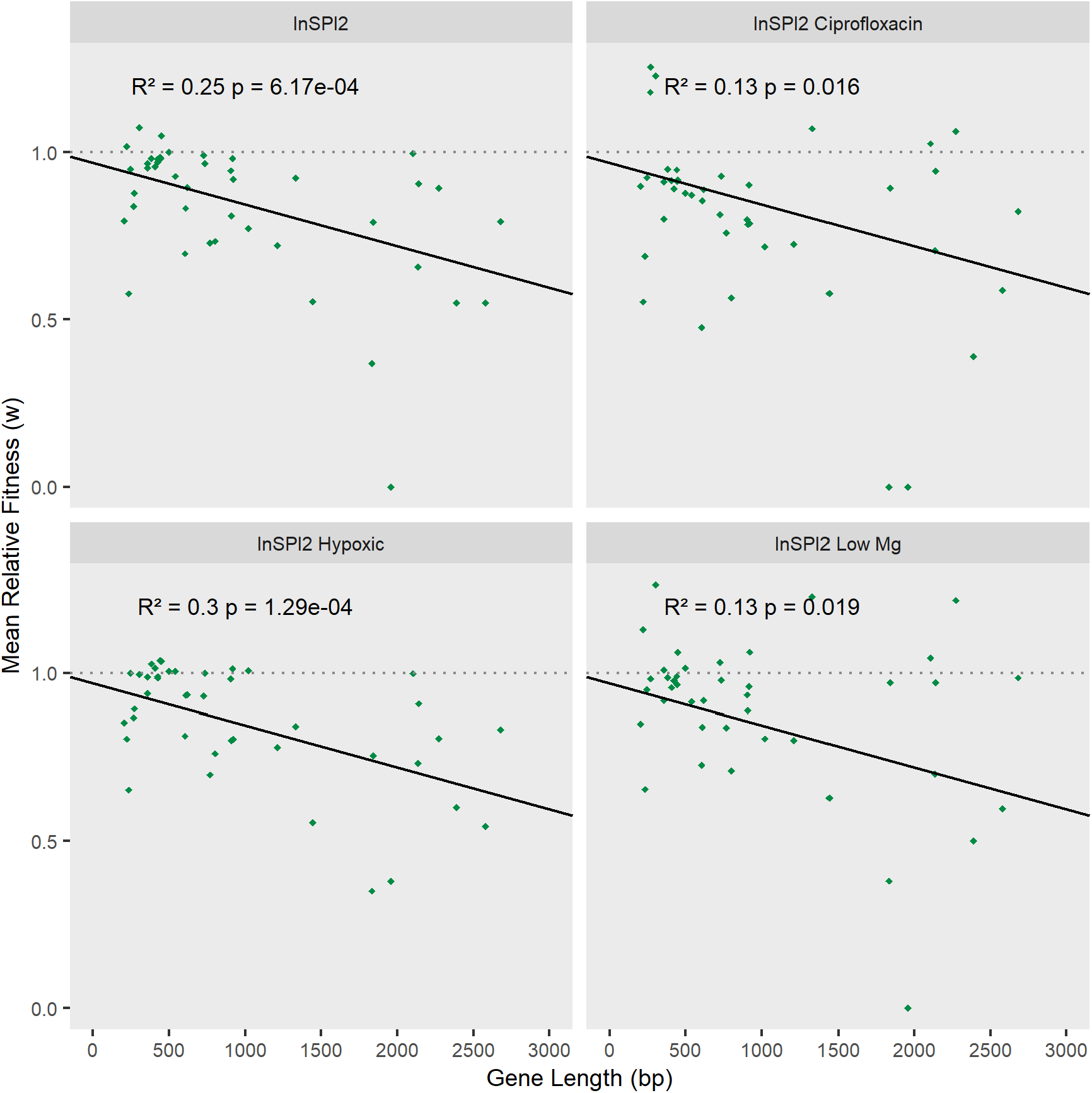
Relationship between relative fitness and gene length. Relative fitness of transferred *<u>E</u>*.*<u>coli</u>* orthologs in S. Typhimurium 4/74 plotted against the gene length in four growth environments. The black line is the regression between the two variables. Grey dotted line shows a fitness of 1. FDR corrected p-values and R^2^ values are shown on the plot.

#### Environment shapes the trajectory of HGT

The impact of the environment on the fitness effects of genes has been well documented in studies of antibiotic resistance genes and metabolic enzymes ^24,57^. Thus, we posed the questions – Do environments affect the central tendency, shape, or spread of the DFEs? Do environments differentially affect the fitness effects of different genes?

The DFEs are fundamental in predicting the response of a population to selection, as the shape of the DFEs can provide an estimation of the effect of a new gene transfer event or mutation ^38^. We observed that certain environments can significantly impact the median (Wilcoxon Rank Sum Test, **Supplementary Table 6 a, Supplementary Figure 4**) and shape or spread of DFEs (Kolmogorov-Smirnov Test, **Supplementary Table 6 b**).

Additionally, we observed a strong interaction between individual genes and the environment (F_126,516_ = 13.32, p < 0.001) suggesting that the environments interact with genes differentially (Figure 7). We observed 29 out of 43 genes showing significant gene by environment (G X E) interactions (see **Supplementary Table 7**). This indicates that the success of HGT events are largely dominated by specific G X E interactions.

**Figure 7.**
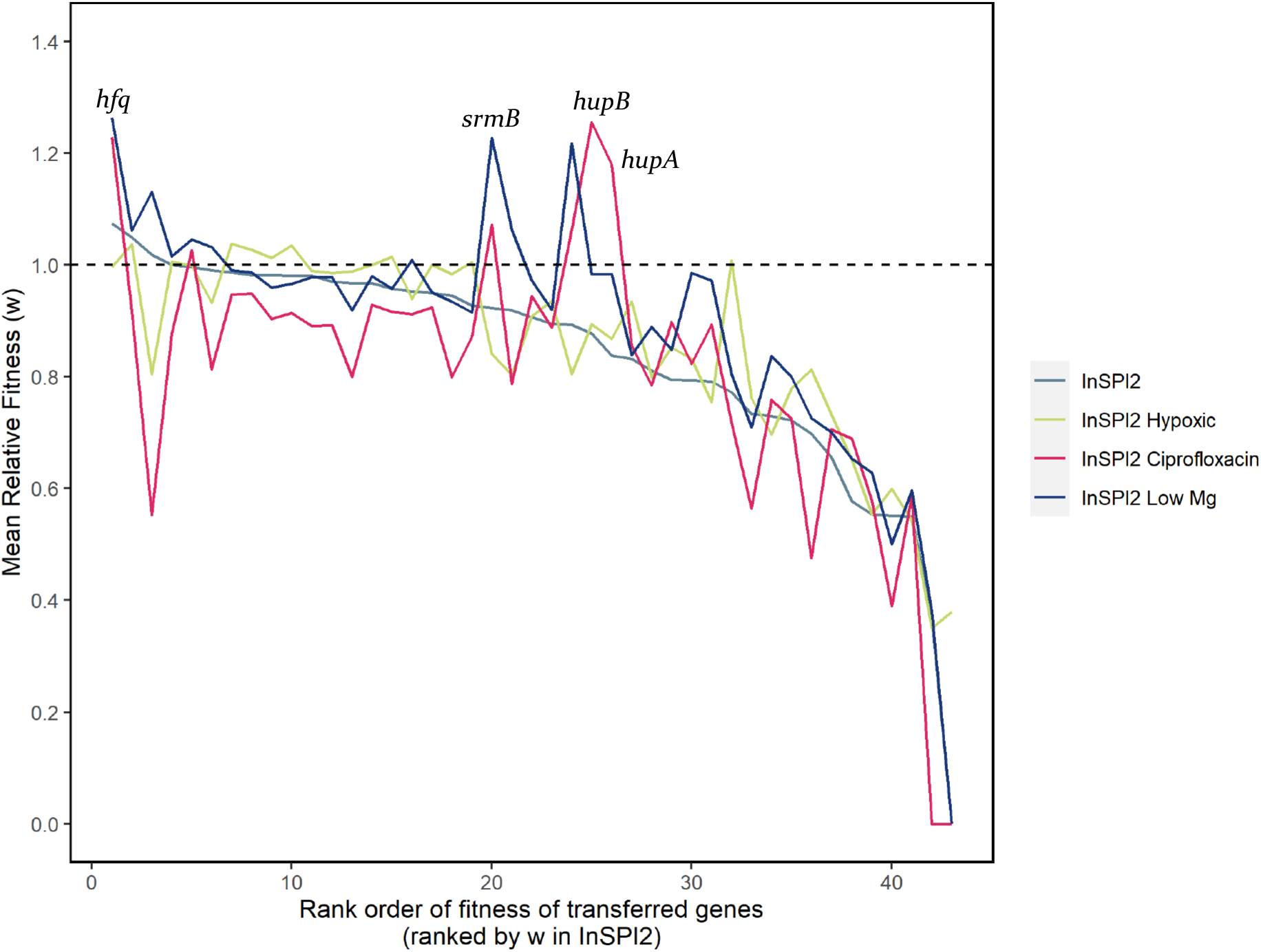
Relative fitness of transferred genes in four environments. The above figure shows the variation in fitness effects of individual genes between environments, few examples have been labelled. Genes are ranked according to the relative fitness measurements in the control environment (InSPI2) only. Black dashed line shows a fitness of 1.

In some cases, the specific function of the gene might be able to explain the observed fitness effect in a certain environment. For example, the *hfq* gene was beneficial for *S*. Typhimurium 4/74 in InSPI2 Ciprofloxacin and InSPI2 Low Mg environments (**Figure 7**). The *hfq* gene is an RNA chaperone facilitating transcriptional regulation in response to environment stress and changes in metabolite concentrations ^58^. Additionally, *hfq* has been shown to regulate the formation of persister cells through the MqsR toxin in *E*.*coli* ^59^. Similarly, *srmB* gene was beneficial for *S*. Typhimurium 4/74 in InSPI2 Ciprofloxacin but had a neutral effect on the host in InSPI2 Low Mg (**Figure 7**). The srmB gene is a DEAD-box RNA helicase facilitating ribosomal assembly ^60^. Additionally, it has been shown that *SrmB* belongs to the AMR Gene Family - ABC-F ATP-binding cassette ribosomal protection proteins; that confer antibiotic resistance via ribosomal protection, unlike other ABC proteins which confer resistance through antibiotic efflux ^61,62,63,64^. However, since ciprofloxacin is known to inhibit DNA replication, the *srmB* gene might have a potential indirect role in regulating fluoroquinolone resistance intracellularly.

#### Protein Dosage is a Barrier to HGT

To elucidate the role of protein dosage, the transferred *E. coli* orthologs were expressed in *S*. Typhimurium 4/74 using different inducer concentrations in each of the growth environments and the fitness for each ortholog gene was compared with the ‘wild type’. We did this by using growth rates as a proxy for fitness. A graphical illustration of high throughput growth curve setup has been provided (Figure 8).

**Figure 8.**
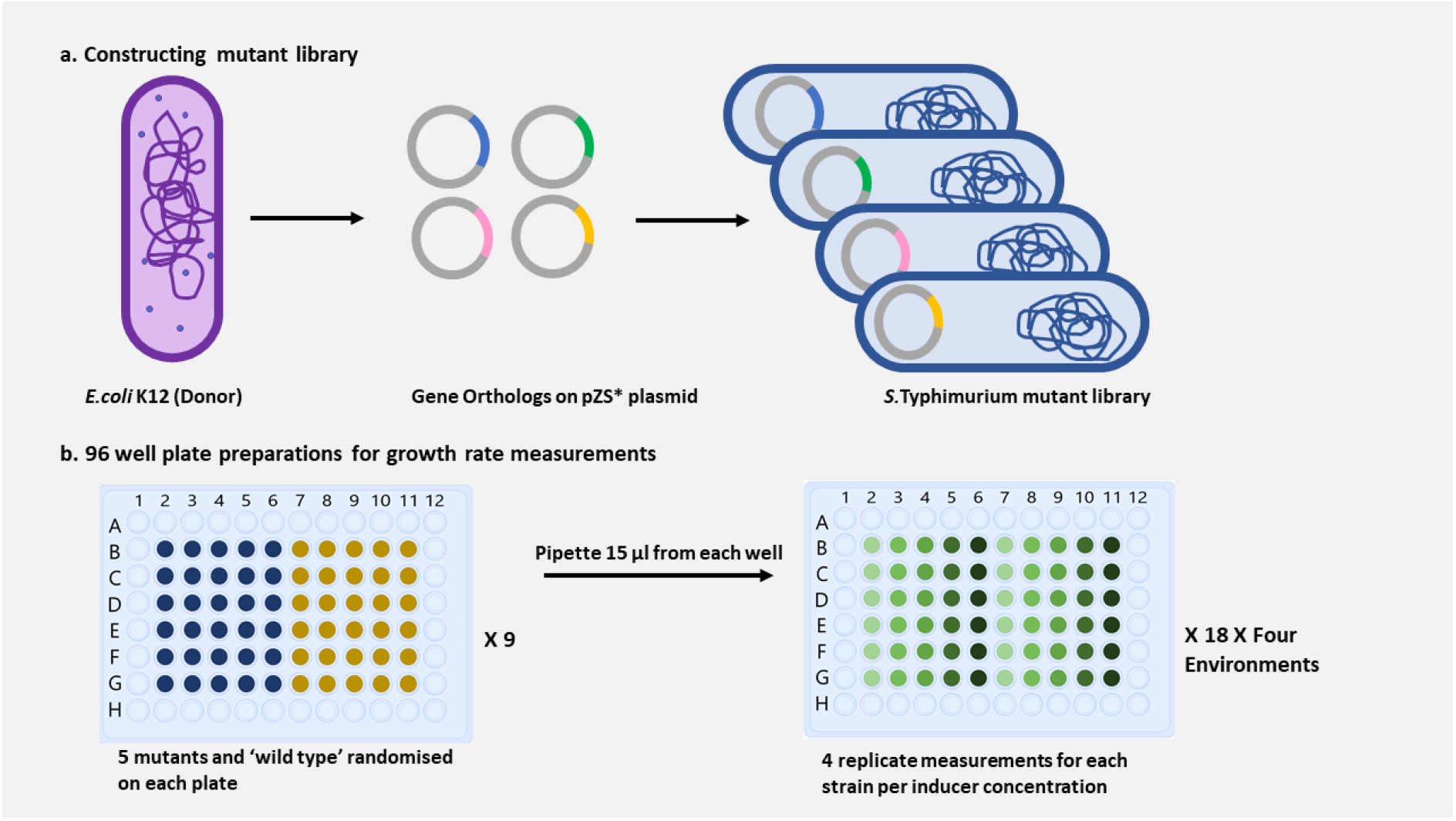
Graphical representation for preparation of mutant library and 96 well plates. **a**. shows the workflow for construction of *S*. Typhimurium strain 4/74 mutant library. **b**. left shows 96 well starter plate preparation, yellow and blue colours showing the two replicates for 5 mutants and a ‘wild type’; and right shows 96 well plate preparation for growth measurements, shades of green from left to right showing increasing inducer concentrations.

Protein dosage is a potential barrier to HGT, as changes in protein concentrations due to variation in gene copy numbers can cause an imbalance in the stoichiometry of the cell, thereby reducing bacterial fitness ^65,66^.

Due to the genetic similarity of *E. coli* and *Salmonella* ^67^, observed fitness effects due to increase in gene expression can be attributed to increased protein levels in the cell (i.e., a dosage effect). To this end, we applied Mandel’s test on the fitness values obtained, and identified 12 genes showing a dosage dependent response (see **Supplementary Table 4**) (**Figure 9**).

**Figure 9.**
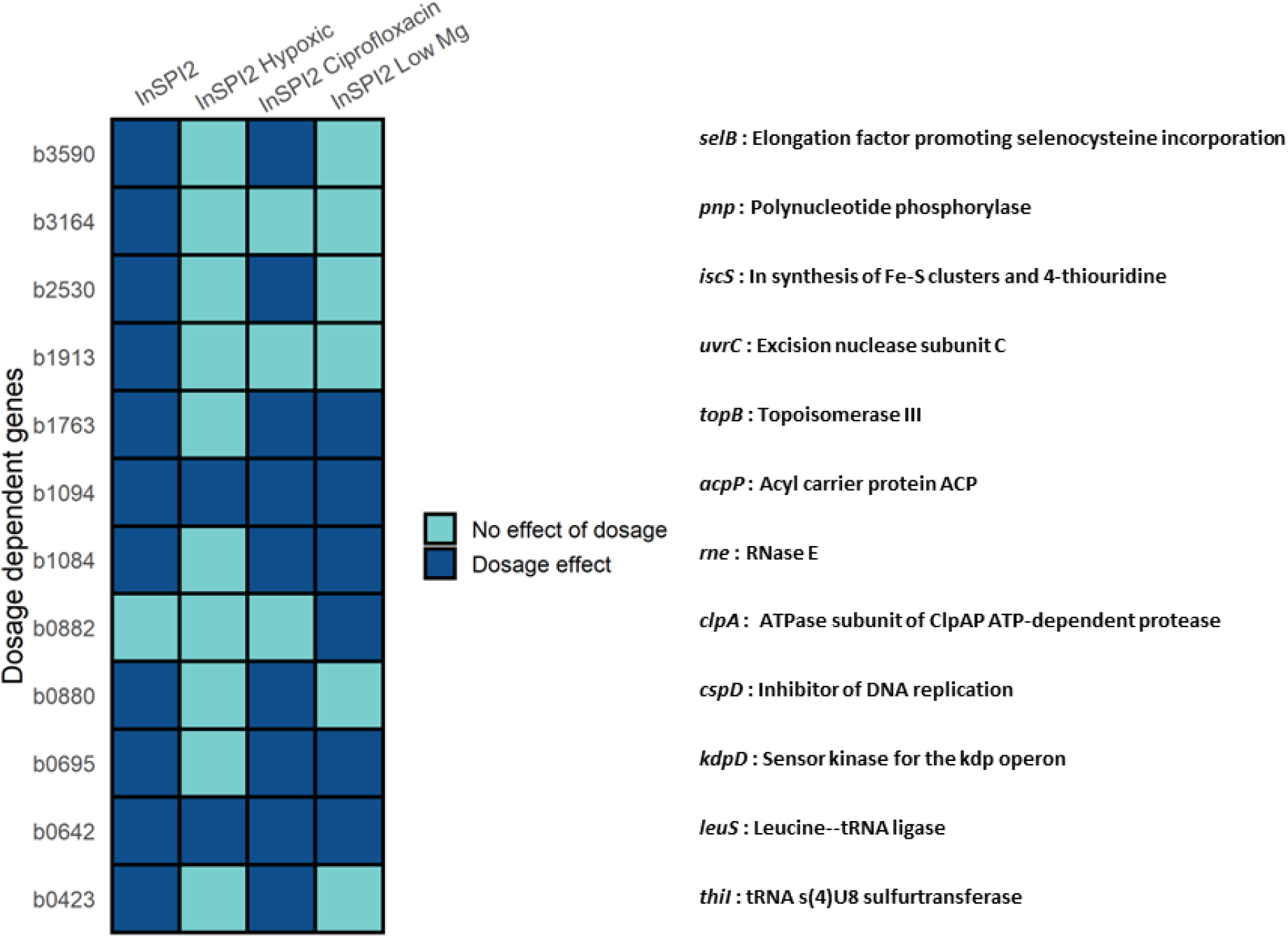
Dosage dependent genes. The figure shows *E*.*coli* orthologs (on y-axis) that exhibit a dosage effect across different environments (on x-axis) when expressed in *S*. Typhimurium 4/74. Gene names and functions are mentioned on the right.

The number of dosage dependent genes were found to be highest in the InSPI2 environment (11 out of 44 genes), followed by InSPI2 Ciprofloxacin (9 out of 44 genes) and InSPI2 Low Mg (6 out of 44 genes), and were lowest in the InSPI2 Hypoxic environment (2 out of 44 genes). In summary, 11 of the 44 transferred genes exhibited dosage dependence in a minimum of two growth environments (**Figure 9**). Classification of all genes based on results from Mandel’s test across all environments are provided as **Supplementary Table 4**.

## Discussion

Despite studies suggesting rampant uptake of foreign genetic material, certain selective forces can act as barriers to HGT ^7,56,68^. This study set out to gain a quantitative understanding of the intrinsic properties of the transferred genes that prevent their successful movement from one species to another. A previous study by Acar Kirit et al. highlighted the importance of conducting experimental fitness measurements in natural environments ^69^, which are scarce due to experimental limitations and their labour-intensive nature. Our study addresses this paucity, by conducting fitness assays in growth environments that capture the conditions encountered by *S*. Typhimurium 4/74 during an intracellular infection ^32^.

Fitness as a measure of reproductive potential is primarily evaluated using bacterial growth rates ^70,71,72^. The density of the microbial culture is directly proportional to the optical density (OD), however, this is only true for a limited OD range, above which measuring culture density with high precision can be demanding ^73^. Additionally, external factors affecting the morphology of bacterial cells, like filamentation, elongation or clumping of cells can interfere with OD measurements, thereby affecting growth rate measurements ^74,75^. Although there were limitations to the use of OD for fitness estimations, the approach allowed us to test fitness effects across different levels of gene expression from no expression to saturation, and to understand the effect of protein dosage in a high throughput and cost effective manner.

Furthermore, we investigated the role of intrinsic properties of the transferred genes and the environment as potential barriers to HGT using NGS. To do this, we pooled the 44 mutants in equal ratios and grew them continuously for ∼8 generations in the four infection relevant environments, and estimated fitness from changes in gene frequencies. The rationale behind pooling of mutants being the approach is time and cost effective, allows for a greater number of replicate experiments, and is less labour intensive. Additionally, studies have shown that copy numbers of a gene are exponential functions of growth rate ^76,77,78^.

Since the aim of the study was to understand what affects the persistence of a newly acquired gene in a population, we tested the immediate fitness effects after a new gene is expressed in the recipient. Although performing pooled growth experiments for a longer duration would help understand the dynamics between mutants due to constrained resources, it could also potentially lead to beneficial compensatory mutations that would balance out the effects of the newly transferred gene. The short duration of the experiment ensured that the energy source was not depleted thereby indicating that the mutants were not competing intensely against each other, and the fitness differences were mainly due to differences in the growth rates of mutants.

Although computational approaches have the benefit of using a large sample size to infer the evolutionary outcomes of HGT, discrepancies in the method of analysis and their underlying assumptions can result in conflicting conclusions. Additionally, these approaches cannot provide information on the role of the environment at the time of the transfer event. This study provides a truly systematic quantitative understanding of the persistence of horizontally transferred genes, and the evolutionary barriers to the emergence and evolution of novel phenotypes. The current study did not observe gene interactions, GC content, and functional category of the genes as significant factors hindering the success of HGT. However, gene length was found to have a significant negative fitness effect on the persistence of horizontally acquired genes. This effect of gene length might be potentially explained by ribosomal sequestration ^79^. An alternate explanation could be the intrinsically disordered protein regions lacking a tertiary structure, which have been found to be correlated with gene length ^69^. These observations are in congruence with the study by Acar Kirit et al. ^69^, suggesting that these barriers are either only weak forces or are below the threshold of practical measurements, especially when the donor and recipient species have diverged recently.

There has been limited support from computational studies for effects of protein dosage as an evolutionary barrier to HGT, mainly because it is difficult to infer dosage/change in protein expression levels from genomic data. To this end, we have measured the fitness effects of the 44 mutants across different levels of gene expression in each of the four infection relevant environments. Our data provides evidence for protein dosage to be a significant barrier to HGT with 12 genes being identified as dosage dependent. Out of the 12 dosage dependent genes, 11 genes were found to be natively expressed at low levels in *S*. Typhimurium 4/74 ^32^. Low levels of native expression potentially suggest that these genes might be maintained by *Salmonella* as single copy and any increase in their protein concentrations potentially imposes a significant fitness cost in the recipient. Interestingly, genes showing a dosage effect were found to be heavily influenced by the environment (Figure 9). Additionally, toxicity due to increased gene expression was also found to be a barrier to HGT (see Supplementary Table 4). A possible mechanism for toxicity effects could be attributed to the poorly optimised gene interactions due to the lack of co-evolution between orthologs or other unknown promiscuous interactions.

Although the importance of the environment has been implicated in the gene acquisition process, we lack an understanding of how the environment determines if the acquired gene is likely to be maintained following HGT. Therefore, analysing the DFE of horizontally transferred genes across different environments is essential to understand if HGT is an opportunistic process or is primarily determined by the intrinsic properties of the newly acquired gene. Our data shows strong G X E interactions, with the fitness effects of the gene becoming unpredictable in each environment. This suggests that improving our knowledge of gene functions could potentially support our understanding of patterns of HGT.

An area of the study that could be expanded upon would be estimating fitness effects of the mutants in macrophages. Even so, the current data represents the closest distribution of fitness effects of horizontally transferred genes achievable using an *in vitro* model. Our data shows which of the environmental variables of a heterogeneous intracellular environment might be favouring HGT; highlighting that the environmental effects should be accounted for when making predictions about the outcome of HGT events. The approach outlined in this study can be further used to study the DFE of genes transferred between phylogenetically distant species to understand which properties of the gene favour or restrict the success of transfer. For example, on transferring orthologs from *E. coli* to *Lactobacillus* certain barriers like GC content or gene interactions might impose significant fitness costs on the recipient thereby hampering the persistence of the newly acquired gene. The fitness costs could arise due to disruption of functional interactions on the introduction of a highly divergent partner or that the divergent genes may be involved in novel negative interactions and that the recipient genome has not evolved to buffer these deleterious side effects.

In conclusion, our data provides evidence that the environment plays a crucial role in influencing the magnitude of gene length and protein dosage as significant evolutionary barriers to the success of gene transfer events.

## Materials and Methods

### Bacterial Strains and Growth Conditions

*Escherichia coli* K-12 MG1655 and *Salmonella enterica* subsp. enterica serovar Typhimurium strain 4/74 ^80^ were used as the donor and recipient strain respectively. LB Broth (Lennox) and SPI-2 inducing PCN (phosphate/carbon/nitrogen) medium (InSPI2 pH 5.8) ^33,34^ were used for growing strains.

### Construction of pZS4-kan-tetR plasmid

Acar Kirit et al. (2020) modified the original pZS4Int plasmid ^35^ by removing the *lacI* gene. The modified pZS4Int plasmid contained the *tetR* gene under the constitutive P_N25_ promoter, spectinomycin resistance gene, and origin of replication pSC101 ^69^. Since the 4/74 strain exhibited growth in spectinomycin concentrations ranging from 80-200μg/ml, the spectinomycin gene on the pZS4Int plasmid was replaced with a kanamycin gene using the NEBuilder HiFi DNA Assembly Reaction Protocol.

### Chromosomal modification of S. Typhimurium strain 4/74

The construct (tetR gene under the constitutive P_N25_ promoter and kanamycin resistance gene) from the pZS4-kan-tetR plasmid was integrated into the lambda phage attachment site of 4/74 chromosome (see **Supplementary Figure 1**) using λ red recombination plasmid pSIM5-tet ^81,82^. The integration site in the modified chromosome of 4/74 strain was Sanger sequenced to ensure no mutations were introduced during the recombineering process.

### Genes under study

The genes of interest to be artificially transferred were selected from a previous study ^69^. *Escherichia coli* strain K-12 substr. MG1655 (GenBank: U00096.2) was used as the donor. Since the study wanted to understand the role of the barriers affecting HGT, genes that are known to be mobile (e.g., phage-related proteins, insertion sequences, transposable elements) were excluded when selecting genes for transfer. Only genes with validated interactions based on LCMS and MALDI after SPA tagging ^83^ were retained, ensuring that the genes were sampled to include a representative range of interactions. Additionally, since a random selection would be biased towards an overrepresentation of genes contained within large functional modules, sampling of the genes ensured that the genes belonged to different functional modules. Furthermore, sampled genes were not accounted for their origins, resulting in 36 genes being part of the core genome, 5 genes predicted to have been horizontally transferred, and 3 genes with unknown origin ^69^.

### Construction of S. Typhimurium 4/74 mutant library

The selected 44 *E*.*coli* orthologs were cloned into a modified version of the pZS* class of plasmids under the control of the p_LtetO-1_ promoter ^35^ and cloned plasmids were electroporated in *E. coli* in a previous study (see **Supplementary Figure 2**). Additionally, a random fragment of the *tetA* gene (721 bp, the mean length of all inserted genes) was cloned without a promoter into the pZS* plasmid and electroporated in E.coli to be used as the ‘wild type’ strain ^69^. A copy of the 45 strains (44 *E*.*coli* genes and a ‘wild type’) were obtained to use in the present study. The insert size was confirmed by PCR on an agarose gel. The plasmids were then electroporated into *S*. Typhimurium 4/74 attλ::tetR-KnR. Successful transformants were selected and after two rounds of streak purification on LB agar plates supplemented with 50 μg/ml Ampicillin, single colonies were grown overnight in liquid LB supplemented with 50 μg/ml Ampicillin and stored at −80°C with 15% glycerol.

### Antibiotic and inducer concentrations

To determine the concentration of ciprofloxacin, growth rate measurements were carried out using Tecan Infinite® 200 Pro Reader. For this, a 1:1000 dilution of an overnight culture of *S*. Typhimurium strain 4/74 attλ::tetR-Kn^R^ carrying the pZS*-tetA (‘wild type’ plasmid) was grown in a 96 well plate in InSPI2 medium supplemented with different ciprofloxacin concentrations and OD_600_ was measured every 20 mins. The concentration of ciprofloxacin to be used in the study was aimed to be twice the doubling time compared to growth in InSPI2.

Gene expression on the plasmids was induced by addition of the inducer anhydrotetracycline (aTc) — a tetracycline derivative with no antibiotic activity — which interacts with TetR to induce gene expression ^84^. To determine the inducer concentration, *S*. Typhimurium strain 4/74 attλ::tetR-Kn^R^ was electroporated with a pZS* plasmid carrying a *gfp* gene under the control of the p_LtetO1_ promoter. Relative Fluorescence Units were measured in InSPI2 with different inducer concentrations using Tecan Infinite® 200 Pro Reader (excitation wavelength, 485 nm; emission wavelength, 535 nm). An arbitrary aTc concentration within the midlinear range of RFU curve was considered as the optimal inducer concentration.

To understand the role of protein dosage as a potential barrier to HGT the transferred genes on the plasmids were induced using increasing inducer concentrations thereby leading to increased protein levels in the cell ^69^. The inducer concentrations used were aTc 0ng/ml (control), aTc 2ng/ml (below optimal expression), aTc 4ng/ml (optimal expression), aTc 6ng/ml (above optimal expression) and aTc 8ng/ml (saturation).

### Growth measurements using Growth Profiler 960

A 1:20 dilution of individual overnight cultures of the 44 *S*. Typhimurium 4/74 attλ::tetR-Kn^R^, pZS* carrying mutants were grown individually in InSPI2 medium supplemented with 50 μg/ml Ampicillin to OD_600_ 0.2. The position of the strains were randomised on 96 well plates to minimise position effects. For each environment, cultivations were carried out in 96-half-deepwell microplates (CR1496dg, Enzyscreen) with gas-permeable sandwich covers and clamps, except InSPI2 Hypoxic, where cultures were overlaid with sterile paraffin oil and gas-permeable sandwich covers were used in conjunction with carbon-flush clamp units. A total volume of 250μl including a 15μl inoculum from the starter plates with randomised layout of genes was used throughout the experiment. The wells were additionally supplemented with different aTc concentrations to induce gene expression (see **Antibiotic and Inducer Concentrations** above). **Figure 8** shows a graphical representation of the methodology. Each experiment was repeated to produce 4 biological replicates across two days. Plates were incubated for 18 hours at 37°C and shaken at 250 rpm, except for InSPI2 Hypoxic, where cultures were grown statically. Measurements were recorded every 20 mins using optimal image scanning by Growth Profiler 960 platform (Enzyscreen). The green values (G-values) obtained for each well were converted into their equivalent OD_600_ values using the image analysis software GP960Viewer (Enzyscreen).

### Pooled Growth Assays

A 1:10 dilution of individual overnight cultures of the 44 S. Typhimurium 4/74 attλ::tetR-Kn^R^, pZS* carrying mutants were grown in InSPI2 medium supplemented with 50 μg/ml Ampicillin to OD_600_ 0.2. The following steps were carried out on ice. After reaching the target OD_600_, strains were mixed at equal volumes to make a total of 135mL. The pooled stock was thoroughly mixed and, 50 aliquots - each containing 1 ml pooled stock and 250μl 80% glycerol, were stored at −80°C. The remaining volume of the pooled stock was split to create two technical replicates for the starting frequency of the pooled growth experiments. To verify the cell concentration of the pooled stock, serial dilutions of a frozen stock were plated on LB agar plates supplemented with 50 μg/ml Ampicillin to obtain CFU counts. The pooled stock concentration was approximately 5 × 10^7^ CFU/ml.

To perform the pooled growth experiments, frozen aliquots of the pooled stock were thawed on ice, and then gently mixed by pipetting. A total volume of 100ml including an inoculum of 107 cells from the pooled stock was used throughout the experiment. Gene expression was induced by the addition of 4ng/ml aTc, and 50 μg/ml Ampicillin was used for maintenance of pZS* plasmids carrying the transferred genes. For each of the four infection relevant environments (**Supplementary Table 1**), 100ml volume of cultures were grown in 200ml conical flasks, except InSPI2 Hypoxic, where cultures were filled up to the brim of 50ml conical tubes, capped with rubber septum, and further sealed using parafilm. The flasks were incubated at 37°C and 250 rpm and grown to OD_600_ 0.4. For the InSPI2 Hypoxic environment, cultures were grown on static at 37°C to OD_600_ 0.2. Four replicate experiments were performed for each of the environments under study. On reaching the target OD (i.e., endpoint of the experiment), plasmid DNA was extracted from the total culture volume using InvitrogenTM PureLinkTM HiPure Plasmid Midiprep Kit (Catalog number: K210004).

### Sequencing

DNA libraries for 16 biological replicates (4 replicate pooled experiments for each of the 4 environments) and 2 technical replicates (starting frequency of pooled growth experiments) were prepared and sequenced on Illumina NovaSeq 6000 platform (PE150bp) by our collaborators at the Laboratories of Molecular Anthropology and Microbiome Research (LMAMR), University of Oklahoma, USA, resulting in ∼3.2 million read pairs per sample.

To briefly describe the library preparation protocol, DNA was first fragmented by sonication using Qsonica 800R, followed by construction for Illumina compatible sequencing libraries using the Kapa Hyperprep kit following manufacturer’s protocols. Sequencing libraries were generated using PCR with Illumina-specific barcoded primers and dual index approach was used to allow multiplexing. The amplified library was then validated using 4150 TapeStation System (Agilent Technologies, Inc.), and size selected using Pippin Prep, BluePippin (Sage Science, Inc.). The libraries were pooled in equimolar concentrations and were sequenced.

### Processing of Sequencing Data

Quality checks and demultiplexing of sequenced reads were performed by our collaborators. Sequenced reads were then processed using AdapterRemoval (v2) ^85^ to trim regions with low quality bases (q<30), merge overlapping read pairs, remove Illumina adapter sequences, and remove reads containing ambiguous bases (Ns).

A customised reference genome consisting of FASTA format sequences of the 44 transferred E. coli orthologs, ‘wild type’ tetA fragment, pZS* plasmid backbone, complete sequences of S. Typhimurium 4/74 chromosome (GenBank: CP002487.1); and its 3 native plasmids TY474p1 (GenBank: CP002488.1), TY474p2 (GenBank: CP002489.1), and TY474p3 (GenBank: CP002490.1) was built using Bowtie2 (Version 2.4.4) (Langmead et al., 2009). The sequenced reads were then mapped to the custom built reference genome using the defaults local alignment parameter in Bowtie2 (Version 2.4.4) (Langmead et al., 2009). The mapped SAM files were converted to BAM format and the corresponding BAM files were sorted using Samtools (Tools for Alignment in SAM format) (Version 1.13) (Li et al., 2009). Read depths at each nucleotide position were calculated using depth function of Samtools (Version 1.13) (Li et al., 2009). Nucleotide depths were converted to gene frequencies by calculating the median of depths obtained for every gene using the R software (Version 4.0.0). All steps of the sequencing data processing were performed on Ubuntu 16.04.5 LTS (GNU/Linux 4.15.0-52-generic x86_64). **Figure 1** provides a graphical illustration of the methodology.

### Intrinsic properties of orthologs and statistical analysis

Effect of protein dosage was studied using growth rates of individual mutants relative to the ‘wild type’. Growth rates for all mutants across all inducer concentrations were estimated from the log linear part of the growth curve through regression analysis. Relative fitness (w) for the transferred genes was calculated by normalising the growth rates of the mutants with the ‘wild type’ for the respective inducer concentrations.

To categorise genes based on dosage effect, Mandel’s test for Linearity was applied to the relative fitness values for every gene to decide if the data points best fit a linear function than a quadratic function. The data used for Mandel’s test is provided as **Supplementary Table 2**. The test assesses if the residual variances of the linear and quadratic model are significantly different using the Fisher– Snedecor F-test ^86,87^.

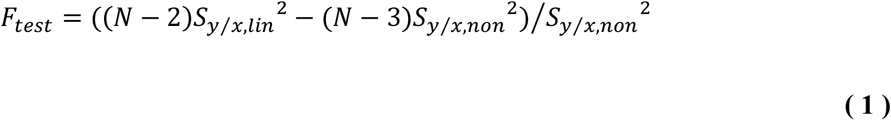

In the above formula, N denotes sample size, S denotes the standard error of regression for the straight line (lin) and for quadratic (non) models.

Other selective factors affecting the DFEs of the transferred genes were studied using the NGS approach. Each replicate pooled growth assay was normalised by the read depth of the vector backbone to account for variation in sequencing library preparation. Fitness costs of selected genes were estimated for each replicate by using the following regression model.

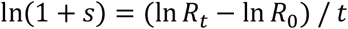

where *R*_0_ and *R*_*t*_ are the ratios of the frequencies of mutant (depth of gene) to ‘wild type’ (depth of a non-functional *tetA* fragment) at start and end point of the experiment, respectively and t is the number of generations ^39^. 43 genes were included in the analysis, as no reads were detected for one of the transferred genes (b1084).

To test whether the genes were neutral or not, a one-tailed one sample t-test was applied on 4 replicate measurements for each gene, with μ_0_ < 1 or μ_0_ > 1. The p-values obtained were corrected for multiple tests using the FDR method ^88^ and α = 0.05 was used as the level of significance.

The transferred orthologs were divided into the two groups using the Database for Clusters of Orthologous Groups (COGs). A one-sided Wilcoxon Rank Sum test was applied on the mean fitness values of genes for 4 replicate measurements to analyse if the fitness effects of informational genes was smaller than those of the operational genes as predicted *a priori* ^12,13,14^.

Additionally, we also looked at other intrinsic gene properties: the number of protein-protein interactions (PPI), metabolic interactions (MI) and regulatory interactions (RI), gene length, GC content and codon usage. The number of protein-protein, metabolic and regulatory interactions for the transferred orthologs in *S*. Typhimurium strain 4/74 were identified using the SalmoNet database (http://salmonet.org/). Gene length was calculated as the number of nucleotides in the coding sequence (CDS) (i.e., from the start to the stop codon) of the transferred *E*.*coli* genes. GC content was calculated as the absolute deviation between the transferred *E*.*coli* genes and their corresponding *S*. Typhimurium strain 4/74 orthologs. Codon usage was estimated as the absolute deviation of the frequency of optimal codon usage (FOP) in the transferred *E*.*coli* genes using the FOP in *S*. Typhimurium (http://www.kazusa.or.jp/codon/cgi-bin/showcodon.cgi?species=602). Multiple Linear regression was used to study the relationship between the intrinsic gene properties and fitness effects, by using the following model.

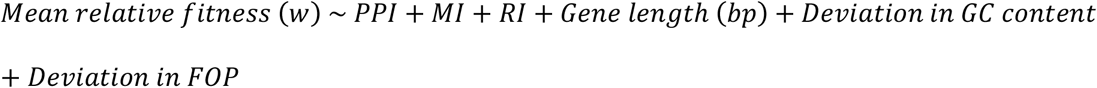

Data used in the model is provided as **Supplementary Table 3**. The p-values obtained were corrected for multiple tests using the FDR method ^88^.

To investigate the effect of the environments on the central tendency of the DFEs, we used a paired and two-sided Wilcoxon Signed Rank Test on each pairwise comparison of the four environments. To test if the shape and spread of the distributions were significantly different between environments, a two-sided Kolmogorov-Smirnov (K-S) test was used. For each of these tests, the fitness values for genes were provided as a mean of 4 replicate measurements. The p-values obtained were corrected for multiple tests using the FDR method ^88^.

The gene by environment (G x E) interactions were studied using a two-way analysis of variance (ANOVA) test for each pairwise comparison for the four environments, using the following model.

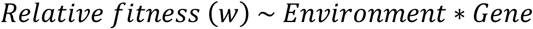

The p-values obtained were corrected for multiple tests using the FDR method ^88^.

All statistical analysis were carried out using the R software (Version 4.0.2) in RStudio (Version 1.1.456).

## Acknowledgements

We thank Wai Yee Fong for sharing S. Typhimurium strain 4/74 and providing guidance on InSPI2 media preparation; Blanca Perez-Sepulveda, Rocío Canals and, Nicolas Wenner for their guidance on chromosomal modification of Salmonella; Simon (Xiaojun) Zhu for his guidance with using Growth Profiler 960; Hawra Al-Ghafli for helping with preparation of pooled stock of mutants; and to, Cecil M. Lewis, Jr., Krithivasan Sankaranarayanan and Hande A. Kirit at LMAMR, University of Oklahoma, USA for performing library preparation and sequencing on our samples. We thank Andrea J. Betancourt for helpful feedback on the manuscript. This work was supported by the European Research Council under the Horizon 2020 Framework Programme (FP/2007-2013) / ERC grant agreement no. [648440] to J.P.B. Library construction and sequencing was supported by OCAST (HF20-032-1) to CML and KS, and National Science Foundation grant (Subaward No. SDSMT-OK18-05) to KS.

## Author Contributions

RPB, HAK and JPB designed the experiments together. RPB performed wet-lab experiments. KS and CML provided materials and support for genome sequencing. KS and HAK constructed genome libraries, sequencing and performed preliminary data quality analysis. RPB processed the sequenced reads with help from HAK. RPB performed statistical analysis. RPB wrote the initial draft of the manuscript and revised it together with HAK and JPB. All the authors have read and approved the manuscript.

