## Supplementary Figures and Tables for "Evolutionary barriers to horizontal gene transfer in macrophage associated *Salmonella*"

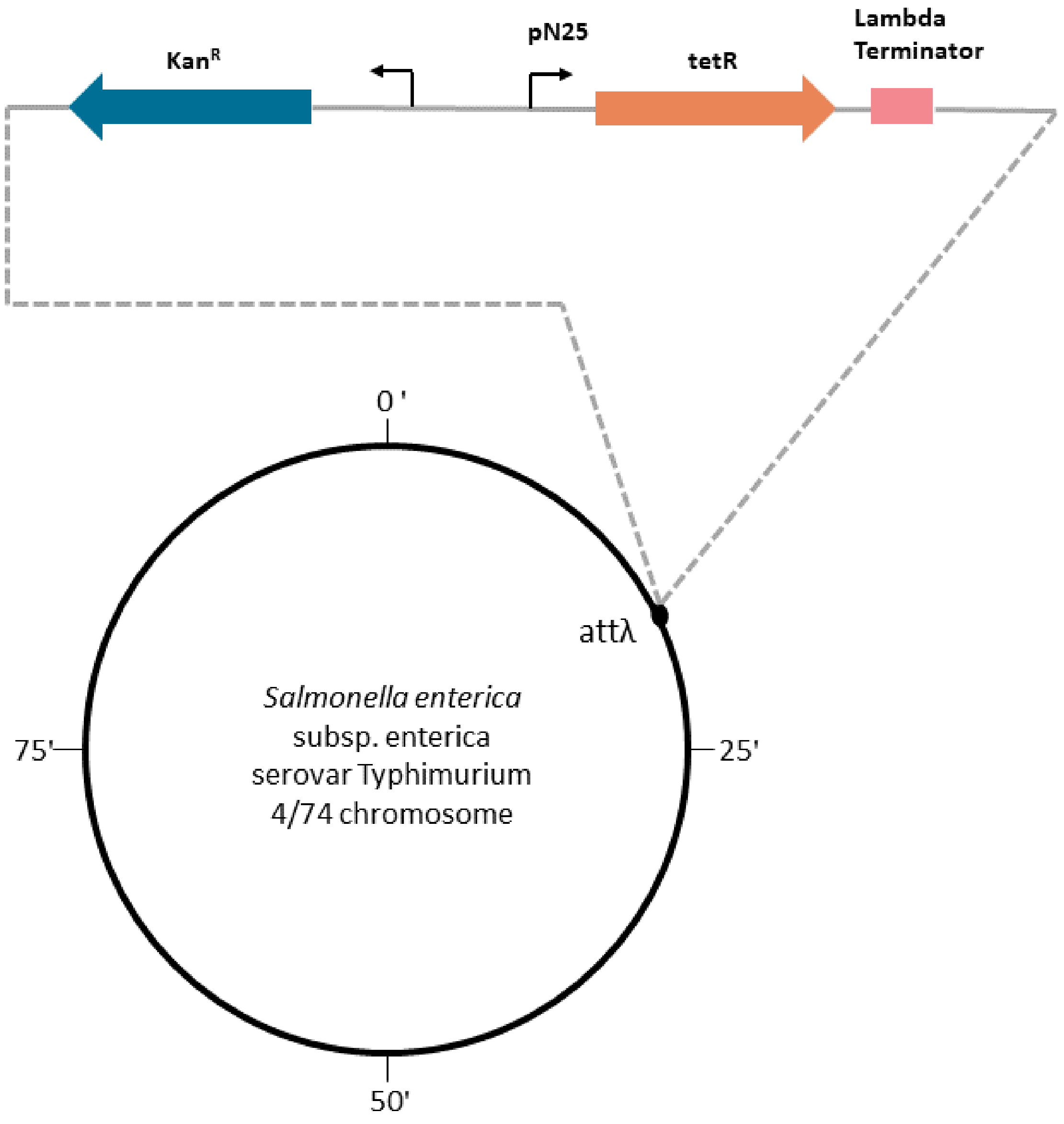


**Supplementary Figure 1 Schematic representation of *Salmonella enterica* serovar Typhimurium attλ::tetR-Kn$^R$ 4/74 strain**

Recipient strain for genes transferred from *Escherichia coli* K-12 MG1655.

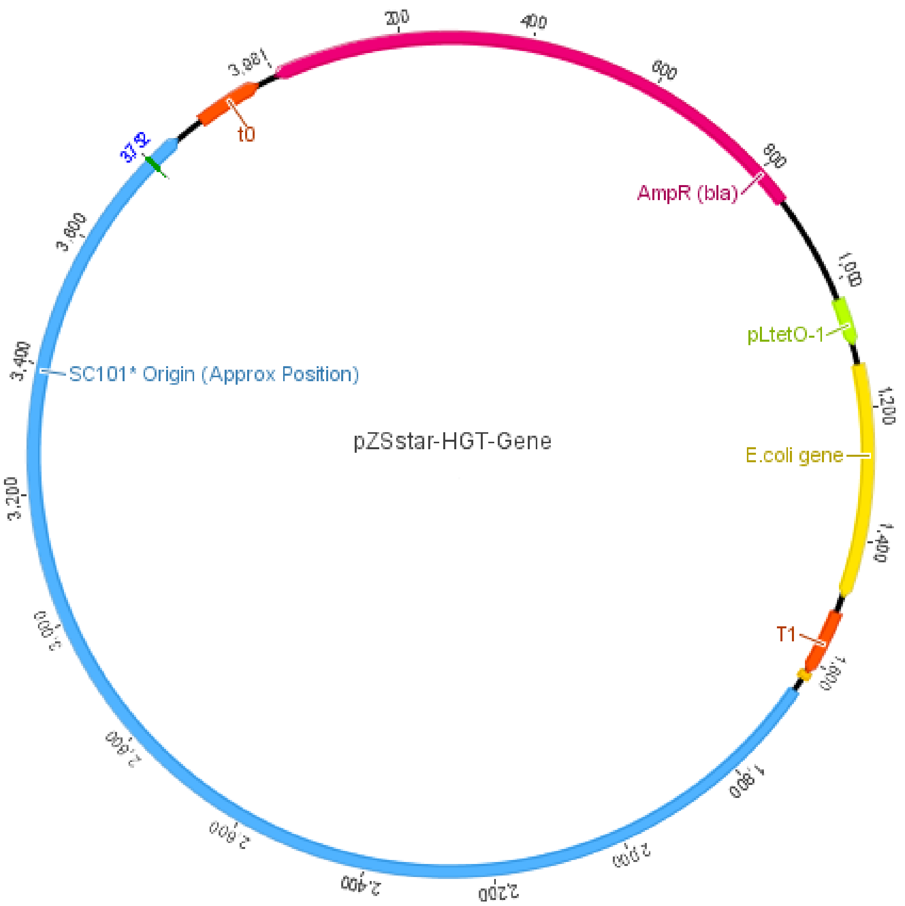


**Supplementary Figure 2 Schematic representation of expression plasmid used to construct 4/74 mutant library**

The pZS* plasmid consisting of an SC101 origin of replication (blue), terminators (t0 and T1)(orange), Ampicillin resistance gene (pink), transferred E.coli gene (yellow) under the control of the inducible pLtetO1 promoter (green).

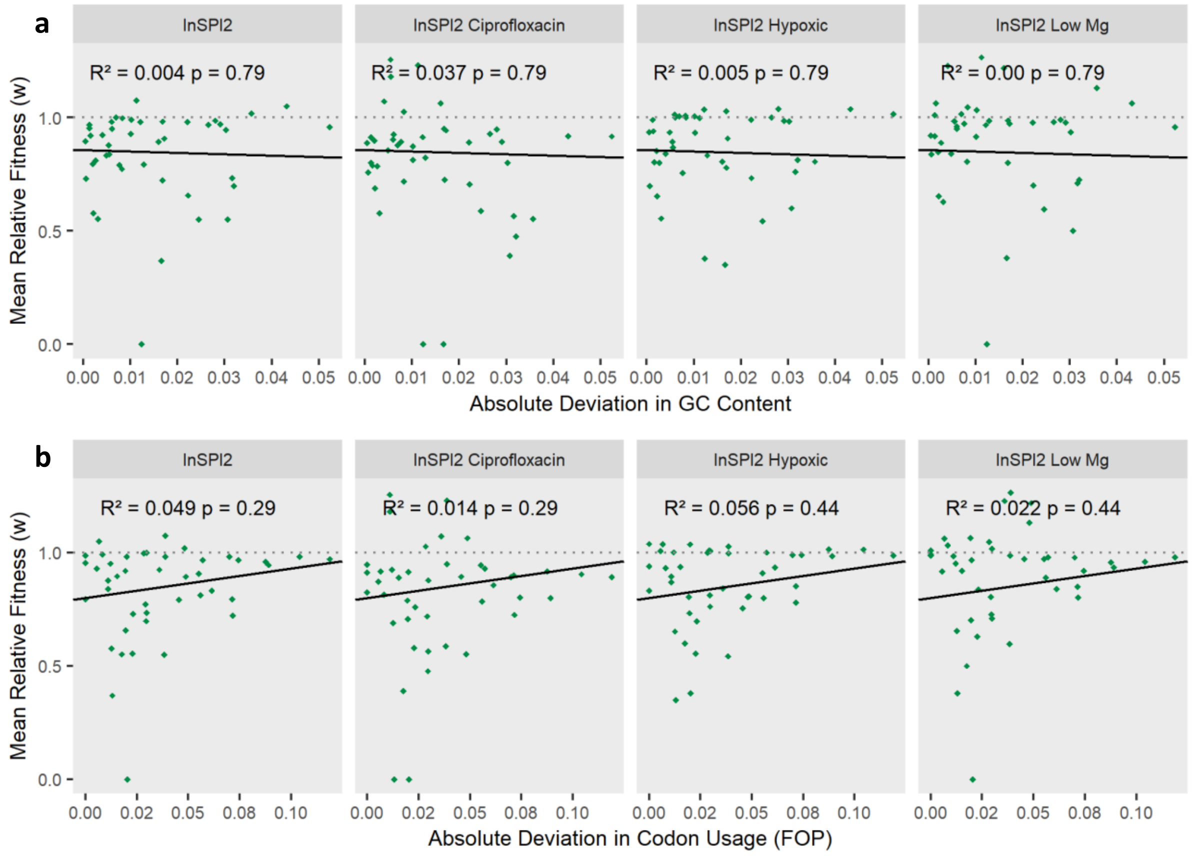


### Supplementary Figure 3 Relationship of GC content and codon usage with fitness

Relative fitness of transferred *E.coli* orthologs in *S.* Typhimurium 4/74 plotted against (a) absolute deviation in GC Content and (b) absolute deviation in FOP (bottom panel) in four growth environments. The black line is the regression between the two variables. Grey dotted line shows a fitness of 1. FDR corrected p-values and $R^2$ values are shown on the plot.

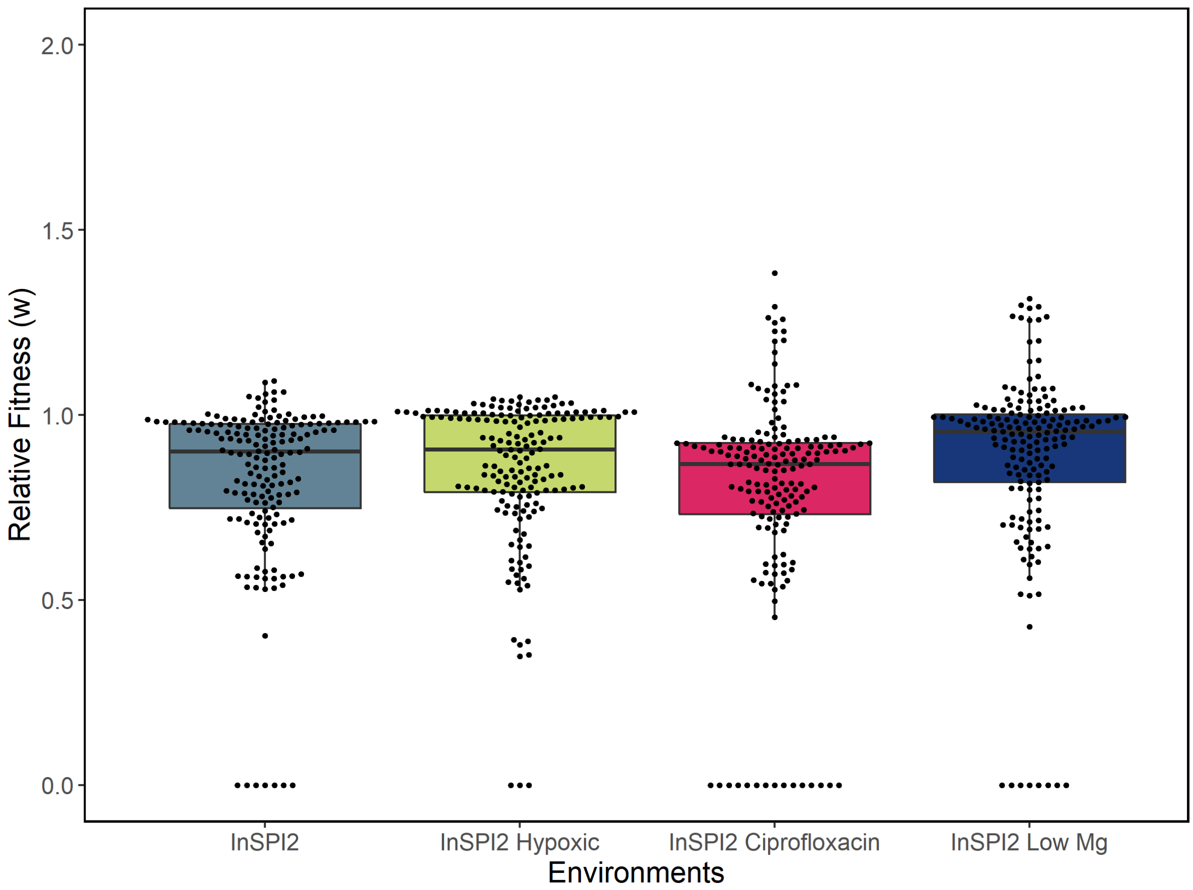


**Supplementary Figure 4 Fitness effects of transferred genes**

Bee swarm boxplot showing the distribution of fitness effects of transferred genes (4 replicate measurements) in four growth environments.

### Supplementary Tables

#### Supplementary Table 1  Infection relevant growth environments

| Growth Environment | Description |
|---|---|
| InSPI2 | PCN medium (pH 5.8) supplemented with 50 µg/ml Ampicillin |
| InSPI2 LowMg$^{2+}$ | PCN (InSPI2) medium with 10 µM MgSO4 supplemented with 50 µg/ml Ampicillin |
| InSPI2 Hypoxic | PCN (InSPI2) medium supplemented with 50 µg/ml Ampicillin and paraffin oil overlay |
| InSPI2 Ciprofloxacin | PCN (InSPI2) medium supplemented with 50 µg/ml Ampicillin and 0.1 µg/ml CIP |

**Supplementary Table 2 – Data used in Mandel's Test (Excel spreadsheet attached).**

**Supplementary Table 3 – Data used in multiple linear regression analysis (Excel spreadsheet attached).**

**Supplementary Table 4 – Transferred *E. coli* orthologs classified into dose response categories based on Mandel's test (Excel spreadsheet attached).**

**Supplementary Table 5 – FDR corrected p-values for fitness effects of *E. coli* orthologs being different from neutral (Excel spreadsheet attached).**

#### Supplementary Table 6a Wilcoxon Signed Rank test

| | InSPI2 | InSPI2 Hypoxic | InSPI2 Ciprofloxacin | InSPI2 Low Mg |
|---|---|---|---|---|
| InSPI2 | _ | p < 0.05 | 0.141 | p < 0.0001 |
| InSPI2 Hypoxic | -2.130 | _ | p < 0.05 | 0.288 |
| InSPI2 Ciprofloxacin | -1.562 | -2.357 | _ | p < 0.0001 |
| InSPI2 Low Mg | -4.176 | -1.061 | -4.363 | _ |

*FDR corrected p-values (upper diagonal) and Z statistic (lower diagonal) for pairwise comparisons using two-sided Wilcoxon Signed Rank test. Comparisons significant with α = 0.05 are shaded.*

**Supplementary Table 6b Kolmogorov-Smirnov test**

| | InSPI2 | InSPI2 Hypoxic | InSPI2 Ciprofloxacin | InSPI2 Low Mg |
|---|---|---|---|---|
| InSPI2 | _ | 0.294 | 0.239 | 0.363 |
| InSPI2 Hypoxic | 0.232 | _ | 0.210 | 0.624 |
| InSPI2 Ciprofloxacin | 0.255 | 0.279 | _ | $p < 0.01$ |
| InSPI2 Low Mg | 0.209 | 0.162 | 0.418 | _ |

*FDR corrected p-values (upper diagonal) and D statistic (lower diagonal) for pairwise comparisons using two-sided Kolmogorov-Smirnov test, $\alpha = 0.05$.*

**Supplementary Table 7 – Genes identified with significant G X E interactions (Excel spreadsheet attached).**
